# Anti-SSTR2 Antibody-Drug Conjugate for Neuroendocrine Cancer Therapy

**DOI:** 10.1101/688184

**Authors:** Yingnan Si, Seulhee Kim, Rachael Guenter, Jianfa Ou, Yun Lu, Kai Chen, John Zhang, Jason Whitt, Angela M. Carter, James A. Bibb, Renata Jaskula-Sztul, James M. Markert, Lufang Zhou, Herbert Chen, Xiaoguang “Margaret” Liu

## Abstract

Neuroendocrine (NE) cancers include a diverse spectrum of hormone-secreting neoplasms that arise from the endocrine and nervous systems. Current chemo- and radio- therapies have marginal curative benefits. This study aimed to develop an innovative antibody-drug conjugate (ADC) to effectively treat NE tumors (NETs). We first confirmed that somatostatin receptor 2 (SSTR2) is an ideal surface target by analyzing 38 patient-derived NET tissues, 33 normal organs, and 3 NET cell lines. We then developed a new monoclonal antibody (mAb, IgG1 and kappa) to target two extracellular domains of SSTR2, which showed strong and specific surface binding to NETs. The ADC was constructed by conjugating the anti-SSTR2 mAb and antimitotic monomethyl auristatin E. *In vitro* evaluations indicated that the ADC can effectively bind, internalize, release payload, and kill NET cells effectively. Finally, the ADC was evaluated *in vivo* using a NET xenografted mouse model to determine cancer targeting, maximal tolerated dosage, pharmacokinetics, and anti-cancer efficacy. The anti-SSTR2 ADC was able to exclusively target and kill NETs with minimal toxicity and high stability *in vivo*. This study demonstrates that the anti-SSTR2 mAb-based ADC has high therapeutic values for NET therapy.

## Introduction

Neuroendocrine (NE) cancers, such as carcinoids, pancreatic islet cell tumors, and medullary thyroid cancer (MTC), arise from cells within the neuroendocrine system that often harbor inherited or sporadic genetic mutations.(1, 2) The United States has an excess of 100,000 NE tumors (NETs) patients with at least 16,000 new diagnoses each year, and there is an estimation of more than 200,000 undiagnosed cases.(3, 4) Patients living with untreatable NET liver metastases have a 5-year survival rate of 13-54%.(5) The fact that 40-95% of patients with NETs are metastatic at the time of initial diagnosis makes complete surgical resections nearly impossible.(3, 6–10) The chemotherapies (e.g., the mTOR inhibitor “everolimus” and the multikinase inhibitor “sunitinib”) have shown limited efficacy and can cause systemic toxicities.(11–18) The somatostatin receptor (SSTR)-targeting analogs (e.g., octreotide and lanreotide) or FDA approved peptide receptor radionuclide therapy (Lutathera^®^) for gastroenteropancreatic NET treatment can extend patients’ life but have a relatively poor response to rapidly proliferating tumors.(19, 20) Thus, it is imperative to develop new treatment strategies.

SSTRs are transmembrane proteins that belong to G-protein coupled receptor (GPCR) family.(21) Many patients overexpress subtypes SSTR1-5 while SSTR2 is predominately found on over 70 NET cells.(22–24) The membrane expression of SSTR2 in NET cells is approximately 20-fold higher than normal cells,(22–24) confirmed by an immunohistochemistry (IHC) analysis performed on a patient tissue microarray (TMA) in this study. Therefore, SSTR2 is a potential target for the development of a new therapeutic approach to treat NETs.

Targeted therapies, such as monoclonal antibodies (mAbs) and antibody-drug conjugates (ADCs) have been applied to treat cancers while minimizing side effects on normal cells.(25–28) ADCs can integrate the advantage of mAbs, such as cancer-specific targeting to minimize side effects, low immunological rejection, long plasma half-life and high stability, and the high cytotoxicity of small molecule chemotherapeutics.(29) After receptor binding, ADC is internalized via receptor-mediated endocytosis, then release the cytotoxic drug into the cytoplasm of cancer cells via either lysosomal degradation or linker cleavage.(30, 31) FDA has approved two ADCs (brentuximab vedotin and trastuzumab emtansine) to treat relapsed Hodgkin lymphoma, systemic anaplastic large cell lymphoma, and relapsed or chemotherapy refractory HER2-positive breast cancer.(32) To our knowledge, neither mAb nor ADC has yet been developed for NET treatment.

The objective of this study was to develop an innovative targeted therapy to treat SSTR2-overexpressing NETs. A surface receptor analysis of multiple patient tissues and normal organ tissues showed that SSTR2 is highly expressed and good target in most NET patients. A new anti-SSTR2 mAb was developed to efficiently target NET and deliver drug via ADC. The specific targeting, maximal tolerated dosage, pharmacokinetics, and anti-cancer efficacy of the anti-SSTR2 ADC were investigated using NET xenografted mouse model. The results showed that the developed ADC was capable of specifically targeting and effectively reducing tumor growth.

## Materials and Methods

The animal studies conform to the Guide for the Care and Use of Laboratory Animals published by the National Institutes of Health and have been approved by the Institutional Biosafety Committee.

### NET patient tissue microarray

The TMA was prepared by Research Pathology Core to analyze the SSTR2 surface expression in NET. The patient tissues were obtained from university Surgical Oncology Tumor Bank through an Institutional Review Board (IRB) approved protocol. The NET microarray consisted of 38 patient tissue cores and 5 normal tissue cores of liver, spleen, placenta, prostate, and tonsil (negative controls). The TMA slides of 33 normal human organs were purchased from US Biomax (Rockville, MD) to confirm the binding specificity of our anti-SSTR2 mAb using IHC staining with NET tissues as positive controls. The tested normal organs are cerebrum, cerebellum, peripheral nerve, adrenal gland, thyroid gland, spleen, thymus, bone marrow, lymph node, tonsil, pancreas, liver, esophagus, stomach, small intestine, colon, lung, salivary, pharynx, kidney, bladder, testis, prostate, penis, ovary, uterine tube, breast, endometrium, cervix, cardiac muscle, skeletal muscle, mesothelium, and skin.

### Cell lines, seed cultures and media

Multiple human NET cell lines, including BON-1 (pancreatic NET), QGP-1 (pancreatic NET), BON-Luc carrying a firefly luciferase reporter gene, MZ-CRC-1 (thyroid NET), and TT (thyroid NET), were used for *in vitro* or *in vivo* studies. BON-1 and MZ-CRC-1 cells were maintained in DMEM/F12 medium supplemented with 10% fetal bovine serum (FBS) in T25 or T75 flasks. TT cells were maintained in RPMI-1640 with 20% FBS. The non-cancerous negative control cell lines, including WI-38 (pulmonary fibroblast) and 917 (foreskin fibroblast), were maintained in DMEM with 10% FBS, 1% non-essential amino acids, and 1% sodium pyruvate. The adherent mAb producing hybridoma was maintained in DMEM with 10% FBS in T flask, while the adapted suspensive hybridoma was cultivated in Hybridoma-SFM with 4 mM L-glutamine and 1% anti-clumping agent (v/v) in shaker flask with agitation of 130 rpm. All seed cultures were incubated at 37 °C and 5% CO_2_ in a humidified incubator (Caron, Marietta, OH). The cell growth, i.e. viable cell density (VCD) and viability, were measured using Countess II automated cell counter or trypan blue (Fisher Scientific, Waltham, MA). All basal media, supplements, and reagents used in this study were purchased from Fisher Scientific or Life Technologies (Part of Fisher) unless otherwise specified.

### Anti-SSTR2 mAb development

Both human SSTR2 (UniProtKB P30874) and mouse SSTR2 (UniProtKB P30875) are an integral membrane glycoprotein with the same topology, including four extracellular topological domains, seven helical transmembrane, and four cytoplasmic topological domains. Protein blast analysis showed that their four extracellular domains had similarity of 81%, 100%, 100%, and 90%, respectively. We developed a SSTR2 mAb to target the 1^st^ extracellular domain (cQTEPYYDLTSNA, aa 33-44) and the 2^nd^ extracellular domain (cALVHWPFGKAICRVV, aa 104-118) using hybridoma technology performed by ProMab. The immune splenocytes with the best anti-SSTR2 antibody expression were fused with myeloma cells (Sp2/0) to obtain 100 hybridoma subclones. The top 4 clones screened using peptides (the 1^st^ and the 2^nd^ extracellular domains)-based ELISA were adapted to serum-free suspensive cultures to produce mAbs.(33) The cancer surface binding of these 4 mAbs was evaluated using flow cytometry and confocal microscopy imaging to define the lead clone which has strong and specific binding to NET (BON-1) cells but low binding to non-cancerous H727 control cells. The isotype of lead clone was determined using mouse antibody isotyping kit (Sigma, St. Louis, MO).

### Anti-SSTR2 mAb production and purification

The mAb production was performed in a 5-L stirred-tank bioreactor controlled at Temp 37 °C, pH 7.0, DO 50% and agitation 70 rpm. The bioreactor was seeded with VCD of 0.3-0.5 ×10^6^ cells/mL in Hybridoma-SFM with 6 g/L glucose, 6 mM L-glutamine, 3.5 g/L Cell Boost #6, and 1% anti-clumping agent. The production cultures were sampled daily to monitor cell growth (i.e., VCD, viability, double time, and growth rate) using cell counter, glucose concentration using glucose analyser, and mAb production using NGC system (Bio-Rad, Hercules, CA). The anti-SSTR2 mAb was purified using our two-step antibody purification protocol by the NGC system (Bio-Rad, Hercules, CA) equipped with Protein A column and ion exchange column.(34, 35)

### ADC construction

Our published cysteine-based conjugation procedure was used to construct ADC in this study. Specifically, the rebridging linker was synthesized to conjugate anti-SSTR2 mAb and MMAE. The generated ADC was purified with PD SpinTrap™ G25 column (GE Healthcare, Chicago, IL) or high-performance liquid chromatography (Waters, Milford, MA). The average drug-antibody ratio (DAR) was calculated as Ratio = (ε_Ab_^248^−Rε_Ab_^280^)/(Rε_D_^280^−ε_D_^248^), where R = A_*248*_/A_*280*_ = Absorbance ratio.(34)

### *In vitro* anti-cancer cytotoxicity (IC_50_)

BON cells were utilized to evaluate the anti-NET cytotoxicity of the anti-SSTR2 ADC and MMAE (control) in 96-well plate following our published protocol.(34) After 3-day incubation, the toxicity was measured through Luminescent Cell Viability Assay (Promega, Madison, MI).

### SDS-PAGE and Western blotting

The Mem-PER plus membrane protein extraction kit was used to extract membrane proteins for surface receptor evaluation. The protein concentration was determined by the Pierce BCA assay. Non-reducing SDS-PAGE was run using NuPAGE™ 4-12% Bis-Tris protein gels. The primary rabbit anti-mouse antibody and HRP-conjugated secondary anti-rabbit antibody were purchased from Abcam (Cambridge, UK). The blotted membrane was treated with Luminata Forte Western HRP substrate (Millipore, Boston, MA), imaged with MyECL imager, and quantified with ImageJ software.

### Flow cytometry

Flow cytometry was performed to quantitate surface receptor binding of SSTR2 mAb using a BD LSRII flow cytometer (BD Biosciences, San Jose, CA). The mAb was labelled with an Alexa Fluor™ 647 labelling kit to generate AF647-mAb. The NET cell lines (BON, TT and MZ) and negative control fibroblast cell line (917) were tested. The detailed procedure was described in literature.(34, 35) The commercial anti-SSTR2 mAb (RD Systems, Minneapolis, MN) was used as control.

### Confocal imaging

Confocal microscope was used to take the imaging of dynamic surface binding and internalization of mAb and ADC in NET cells following our established protocol.(34, 35) Specifically, BacMam GFP Transduction Control was used to stain the cytoplasma and nucleus, and the AF647-mAb or AF647-ADC was used to target cells. The stained cells were observed using Olympus 1X-81 confocal microscope with Olympus FV-1000 laser scan head using a confocal microscope (Olympus IX81, Center Valley, PA). The MitoSox images were recorded and analyzed offline with the ImageJ software.

### Pharmacokinetics study

To investigate the metabolic rate of ADC, 5 different dosages (4, 8, 12, 16, 20 mg/kg-BW) of ADC were injected to 5 groups of randomized mice. Blood samples were collected from tails at 2, 5, 24, 48, 72, 120 hrs post-injection (6 time points in total for each mouse). Blood was centrifuged at 2,000 g for 5 mins to precipitate cells and the supernatant was collected for ELISA analysis. The recommended dose (D) and dosing interval (τ) were calculated as D = C_max(desired)_ · k_e_·V_d_·T·(1−e^−keτ^)/(1−e^−keT^) and τ = ln(C_max(desired)_/C_min(desired)_)/k_e_ + T using previously developed PK model.(36) The calculated D was used in the anti-cancer efficacy animal study.

### *In vivo* anti-NET efficacy study

The NET xenograft mice with tumor volume of 50-60 mm^3^ were randomized to 3 groups (*n* = 6): saline, anti-SSTR2 mAb, and mAb-MMAE conjugate. The mAb or ADC was administrated through tail vein following a dose of 8 mg/kg-BW (determined from PK study) in 50 μL. The same volume of mAb or saline was injected in control groups. The tumor volume and mouse body weight were measured every two days. Four injections were conducted with average injection interval of 4.5 days during the entire treatment period. In the end of experiment, mice were sacrificed to collect tumors and other organs (e.g. brain and liver) for further analysis.

### Hematoxylin and eosin (H&E) staining

The section was deparaffinized before staining and incubated with 200 μL of hematoxylin solution for 5 mins at 25 °C. The dye was washed using tap water and the section was rinsed in PBS for 5 mins. Then, the section was stained in 400 μL of eosin Y solution for 30 seconds and dehydrated twice in absolute alcohols for 2 mins and cleared in xylene.

### Immunohistochemistry staining

Tissue microarray slides were rehydrated using xylene and ethanol, then immersed in citrate buffer (BioGenex, Fermont, CA) for a 10-min pressure cooker cycle to achieve antigen retrieval. Endogenous peroxidase activity was quenched by incubating slides in 3% hydrogen peroxide for 10 mins. Blocking was performed for 1 hr at RT using 3% goat serum and 0.3% Triton-X100 in PBS. SSTR2 was detected with an overnight 4 °C incubation using 1.8 mg/mL of anti-SSTR2 mAb with final concentration of 10 μg/mL, followed by an anti-mouse biotin-labeled secondary antibody and HRP streptavidin. Slides were stained with DAB Chromogen (Dako Liquid DAB+ substrate K3468) and counter stained with hematoxylin. Before being cover slipped and imaged, slides were dehydrated and cleared using ethanol and xylene.

### Statistical analysis

All the data were presented as mean ± standard error of the mean (SEM). Two-tailed Student’s *t* tests were used to determine the significance between two groups. Comparison among multiple groups was performed using a one-way ANOVA followed by post-hoc (Dunnett’s) analysis. The sample size of animal study was determined by our prior study and published ADC therapy study.(37) Statistical significance with \*\*\**P* value of < 0.001 was considered for all tests.

## Results

### SSTR2 overexpression in NET but not normal organs

NET tissue microarray slide was first stained with H&E to confirm the presence and location of NET cells in each core (Fig. 1A), then applied with IHC staining to evaluate the surface expression of SSTR2. The IHC staining demonstrates that approximately 71% of the patient tissue cores are positive for SSTR2 with strong cell membrane localization (Fig. 1B), but SSTR2 was not detectable in normal liver, spleen, placenta, prostate, and tonsil tissue cores (negative controls). The IHC staining of the 33 types of normal human tissues with our anti-SSTR2 mAb showed that there was no detectable SSTR2 surface expression in most of these normal organs except pancreas and skin which had weak signal (Fig. 2A) while the 10 NET tissues (positive controls) had strong signal. The Human Atlas Project database reported a high level of SSTR2 mRNA in brain, lung, liver, muscles, skin, placenta, prostate, tonsil, and pancreas, but the high-resolution images of these normal organs only showed minimal or undetectable surface SSTR2 receptor (Fig. 2B). In addition to tissue microarray analysis, Western blotting also showed a high-level expression of SSTR2 in two pancreatic NET cell lines (BON-1 and QGP-1) and a pulmonary NET cell line (H727), but there was minimal expression in non-cancerous, fibroblast cell lines (917 and WI-38) (data not shown).

**Figure 1.**
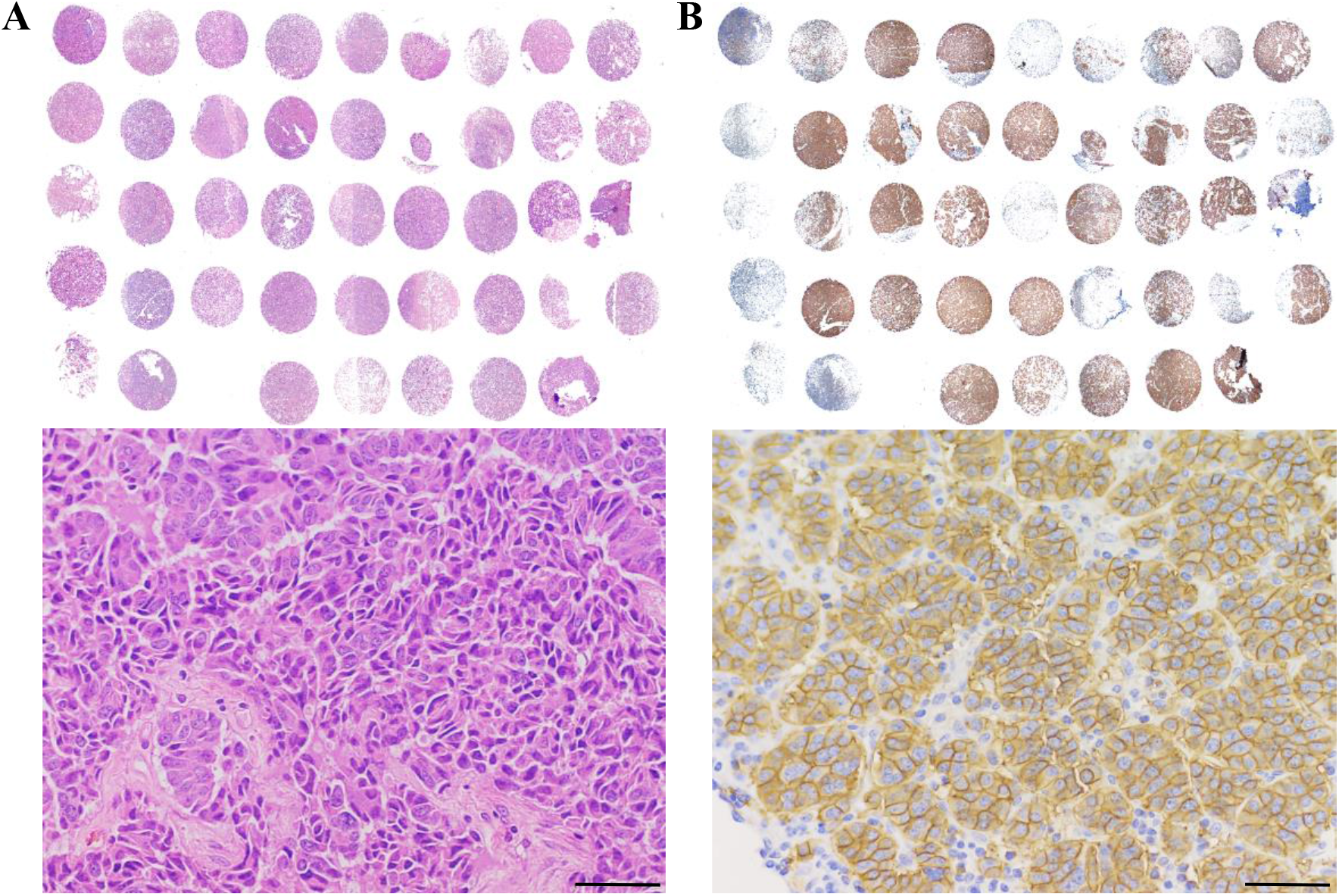
Tissue microarray (TMA) to detect SSTR2 expression in patients. **A,** H&E staining of the TMA including human pancreatic NET tissues (columns 2-9, *n* = 38) and normal tissues (control, column 1, *n* = 5). **B,** IHC analysis of SSTR2 in the TMA. Scale bar equals to 20 μm.

**Figure 2.**
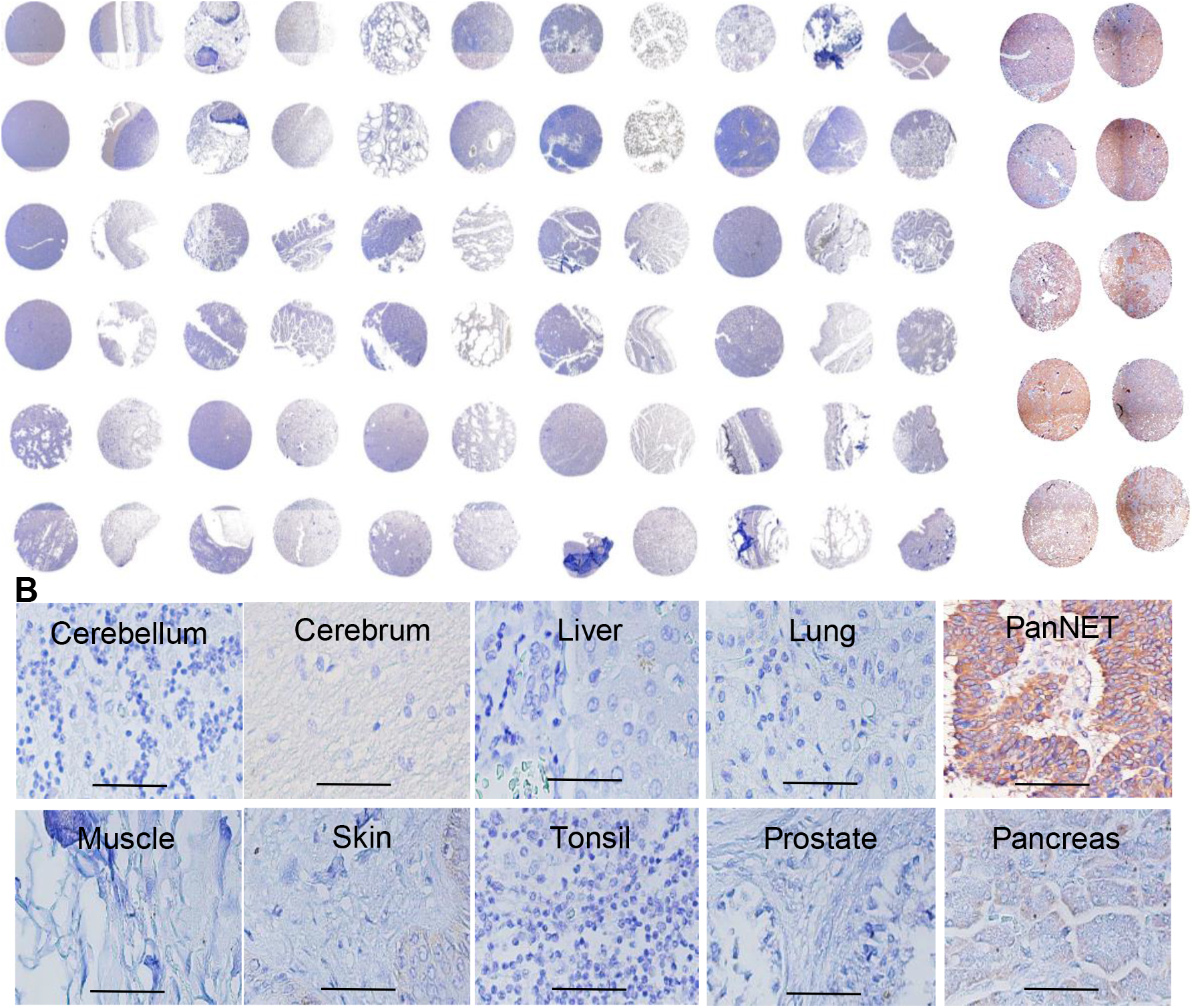
Evaluation of the NET-specific targeting of our anti-SSTR2 antibody using normal human organs or NET tissues by IHC. **A1,** Surface SSTR2 staining in 33 normal human organs (US Biomax, FDA662a, *n* = 2), including cerebrum, cerebellum, peripheral nerve, adrenal gland, thyroid gland, spleen, thymus, bone marrow, lymph node, tonsil, pancreas, liver, esophagus, stomach, small intestine, colon, lung, salivary, pharynx, kidney, bladder, testis, prostate, penis, ovary, uterine tube, breast, endometrium, cervix, cardiac muscle, skeletal muscle, mesothelium, and skin. **A2,** SSTR2 staining on the cell surface in pancreatic NET patient tissues (*n* = 12). **B,** Representative high-resolution IHC imaging of cerebellum, cerebrum, liver, lung, muscle, skin, tonsil, prostate, pancreas, and pancreatic NET. Scale bar equals to 50 μm.

### Development of anti-SSTR2 mAb to target NET

The hybridoma clones secreting anti-SSTR2 mAb were screened using ELISA to identify the top 4 mAb clones that have high binding to the 1^st^, the 2^nd^ or both extracellular domains of SSTR2 (Fig. 3A). In flow cytometry analysis, the surface binding capacity of these 4 mAbs to BON-1 cells was 50%, 80%, 90% and 98%, respectively (Fig. 3B). The clone 4 was defined as “lead clone”, fully characterized, and used throughout the remainder of this study. An isotype analysis revealed that the lead clone is IgG1 kappa, and SDS-PAGE analysis confirmed its molecular weight of 150 kDa (Fig. 3C). Further evaluation showed that the anti-SSTR2 mAb had high surface binding to NET cell lines BON-1 and QGP-1 (>90%) and low binding to fibroblast cell lines 917 and WI-38 (<7.5%) (Fig. 3D). Additionally, we and GenScript isolated, cloned and sequenced the mAb, and confirmed the novelty of our anti-SSTR2 mAb (PCT patent TH Docket No. 222119-8030). To optimally produce mAb, we adapted the adherent hybridoma cells to suspensive culture in stirred-tank bioreactor (Fig. 3E). The cultures in T-flask, spinner flask, and stirred-tank bioreactor generated 8.6, 39.8, and 53.3 mg/L of anti-SSTR2 mAb with a specific growth rate of 0.016, 0.024 and 0.035 hr^−1^, respectively (Fig. 3F).

**Figure 3.**
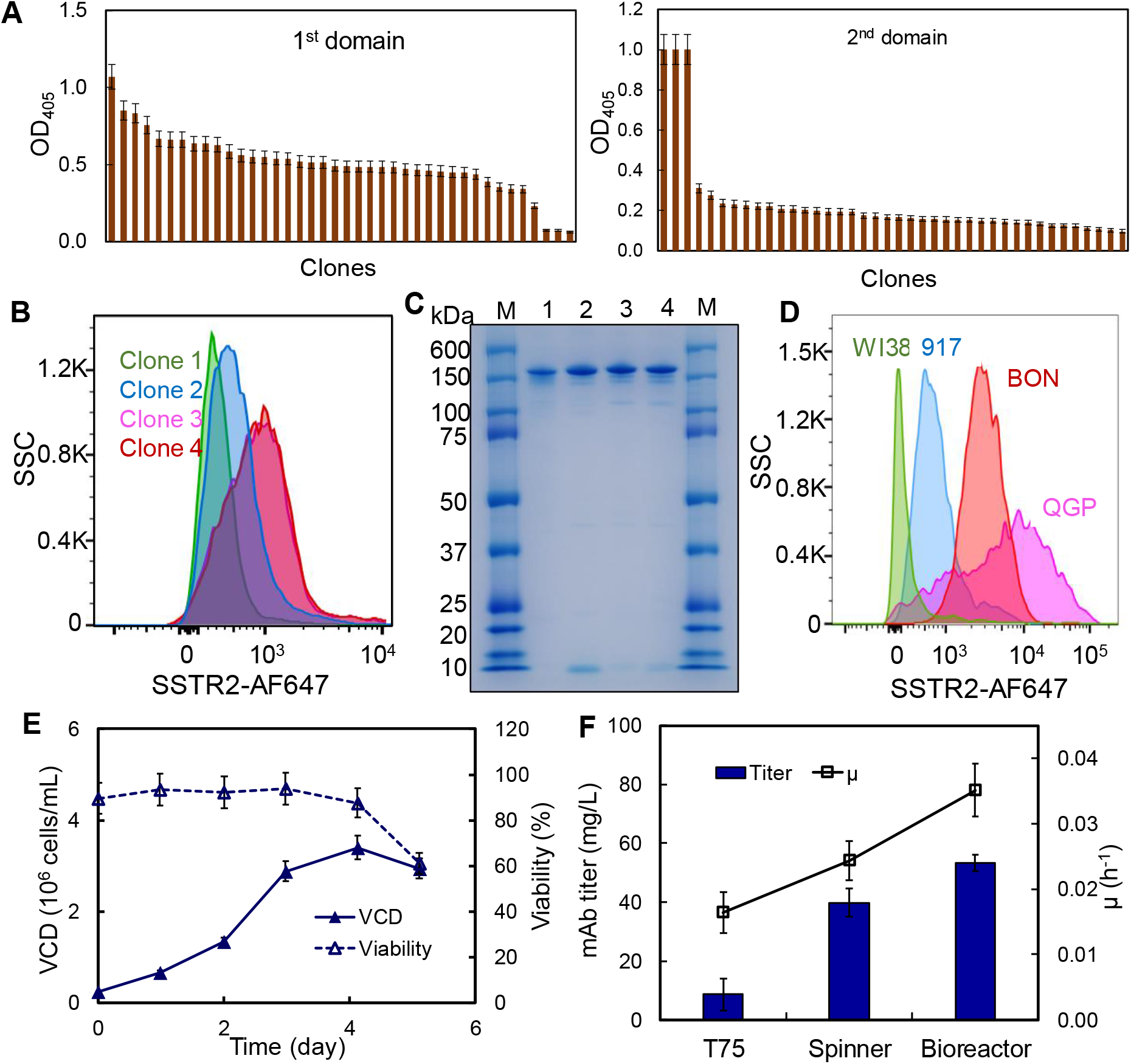
Anti-SSTR2 mAb development and production. **A,** Rank of top anti-SSTR2 mAb clones based on the titer in ELISA screening (data represent mean ± SEM, *n* = 3). **B,** Evaluation of top 4 clones using flow cytometry. **C,** SDS-PAGE to confirm the integrity and purity of mAb (M: marker; 1-4: Clones 1-4). **D,** Evaluation of SSTR2 binding of lead clone in control cell lines (WI38 and 917) and NET cell lines (BON and QGP). **E,** mAb production and hybridoma cell growth in fed-batch suspension cultures (data represent mean ± SEM, *n* = 3). Viable cell density (VCD): 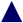, cell viability: 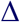, specific growth rate (μ): 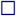.

### High surface binding to NET

To assess the *in vitro* NET-specific targeting of our anti-SSTR2 mAb, we performed live-cell, dynamic CLSM imaging and flow cytometry analysis. As shown in Fig. 4A, the AF647-mAb accumulated on the BON-1 cell surface, displayed as a “red circle”, within 20 mins post incubation due to immunoaffinity. The mAb was then internalized through endocytosis and localized in cytoplasm (detected with BacMam GFP control) within 40 mins. Fig. 4B showed that our anti-SSTR2 mAb had much stronger surface binding to BON-1 cells than the commercial mAb (R&D Systems), 95% vs. 38%.

**Figure 4.**
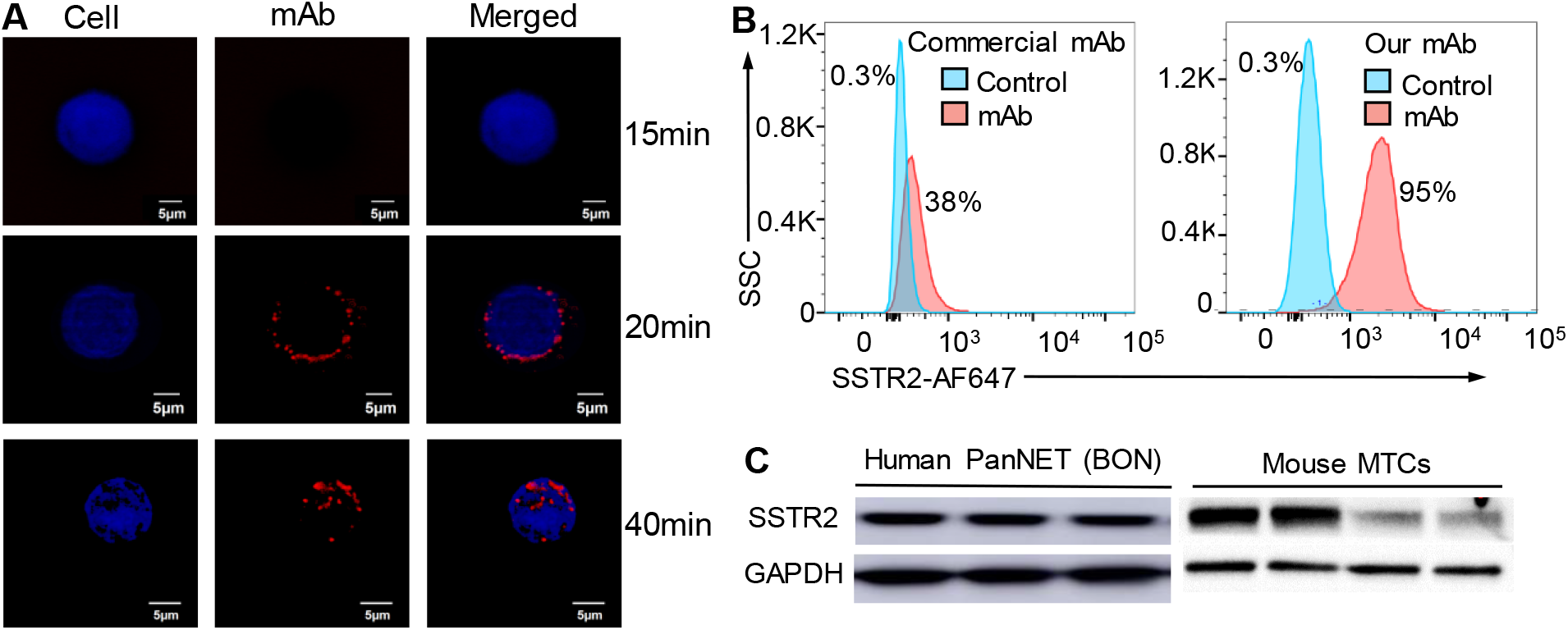
Evaluation of surface binding by our anti-SSTR2 mAb. **A,** Live-cell CLSM dynamic imaging of anti-SSTR2 mAb. Two-color CLSM: whole cell labeled with GFP (displayed as blue) and SSTR2 mAb-MMAE labeled with AF647 (red). Scale bar equals to 5 μm. **B,** Flow cytometry to analyze the surface binding of our anti-SSTR2 mAb to NET cell (BON-1) and negative control cell (917). Stained with 1 μg of mAb-AF647/million cells on ice for 30 mins. **C,** Western blotting of human NET (BON) xenografted tissue and mouse MTC tissues (*n* = 3-4) using our mAb.

The UniProtKB database shows that human SSTR2 (UniProt P30874) and mouse SSTR2 (UniProt P30875) have the same topology, and the 1^st^ and the 2^nd^ extracellular domains of human SSTR2 that our mAb targets have 100% similarity with mouse SSTR2. The Western blotting in Fig. 4C confirmed that our anti-SSTR2 mAb can detect the SSTR2 present in human BON-1 xenografts and in isolated medullary thyroid carcinoma (MTC) tissue from a spontaneous MTC mouse model.(38, 39)

### Anti-SSTR2 ADC construction

We employed our cysteine-based conjugation procedure to construct ADC.(34) Herein, the rebridging peptide-based linker was synthesized to maintain high integrity of the mAb (Fig. 5A) during the conjugation with MMAE. The Agilent 6500 Q-TOF LC/MS confirmed the right structure of linker (Fig. 5B), and SDS-PAGE confirmed the high integrity of ADC structure (Fig. 5C). The DAR of the constructed ADC was approximately 4.0.

**Figure 5.**
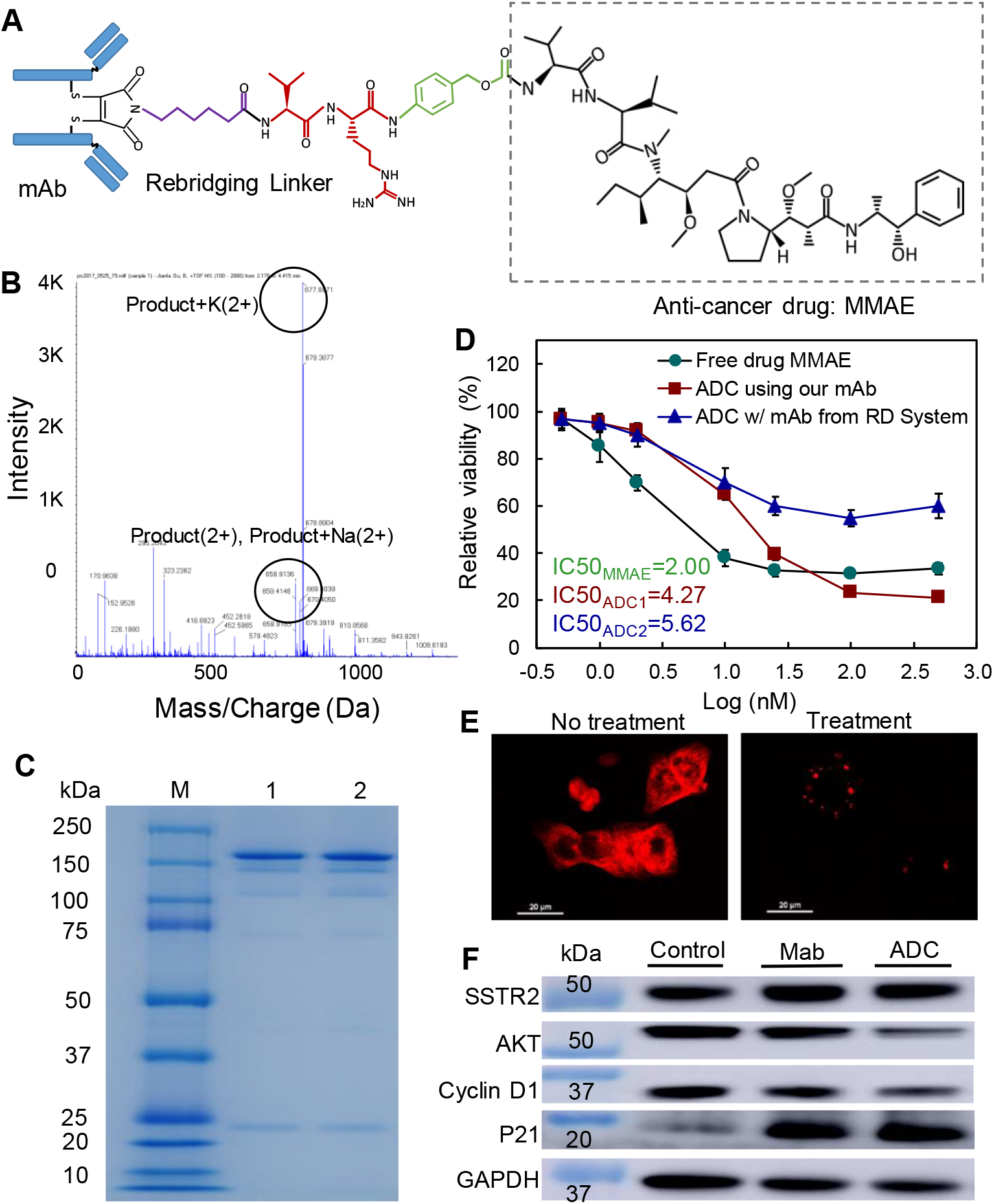
ADC construction and *in vitro* characterization. **A,** Molecule structure of anti-SSTR2 mAb-MMAE using re-bridging linker. **B,** MS analysis to confirm the right structure and proper conjugation of linker-MMAE drug. **C,** The IC_50_ anti-cancer toxicity of free drug 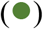, ADC constructed using commercial anti-SSTR2 mAb (R&D Systems, 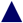), and ADC constructed using our anti-SSTR2 mAb 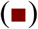 (data represent mean ± SEM, *n* = 3). **D,** SDS-PAGE to check the integrity of mAb-MMAE. **E,** Microtubule de-polymerization in BON cell line treated with MMAE. Scale bar equals to 20 μm. **F,** Western blotting to analyze other anti-cancer mechanisms.

### *In vitro* anti-cancer toxicity of ADC

We evaluated the *in vitro* anti-cancer toxicity of the ADC in BON-1 cells by comparing free drug (MMAE) and two ADCs constructed using the anti-SSTR2 mAb developed in this study or the R&D Systems mAb. MMAE is a highly potent cytotoxin that can block microtubulin polymerization but has never been tested in NETs.(40–43) In this study, the IC_50_ values of MMAE, ADC from our mAb, and ADC from the commercial mAb were 2.00, 4.27, and 5.62 nM, respectively (Fig. 5D). It is clear that the mAb-MMAE ADC had similar nanomolar cytotoxicity to NET cells as free drug.

### Anti-cancer mechanisms

As shown in Fig. 5E, the free drug released from ADC inhibited NET cell proliferation via microtubule de-polymerization. To discover other potential anti-cancer mechanisms of ADC, we analyzed several markers associated with cell proliferation signaling pathways in the BON-1 cells treated with ADC for three days. Western blot showed that both anti-SSTR2 mAb and ADC could block cell proliferation signaling via the PI3K-AKT pathway, downregulate the oncogene Cyclin D1, and induce cell cycle arrest as detected by marker p21 (Fig. 5F).

### Maximum tolerated dose (MTD)

To investigate the MTD, 5 different doses of anti-SSTR2 ADC were injected into Nude (nu/nu) mice (non-tumor bearing) via tail vein: 4, 8, 12, 16, and 20 mg/kg BW (*n* = 2). Mice were monitored twice daily for a total of 21 days and showed no signs of behavior changes such as water intake, labored breathing, rapid weight loss, or impaired ambulation. As shown in Fig. 6A, ADC at a dose range of 4-20 mg/kg BW had no obvious side effects on mice body weight or overall survival. In the end of study, mice were sacrificed and major organs (brain, lung, heart, kidney and liver) were collected for further studies. As shown in H&E staining (Fig. 6B), the brain tissue (and other tissues) had no obvious morphology change, inflammation or apoptotic after ADC treatment. These results indicated that the anti-SSTR2 ADC therapy had no evident off-target effects *in vivo*.

**Figure 6.**
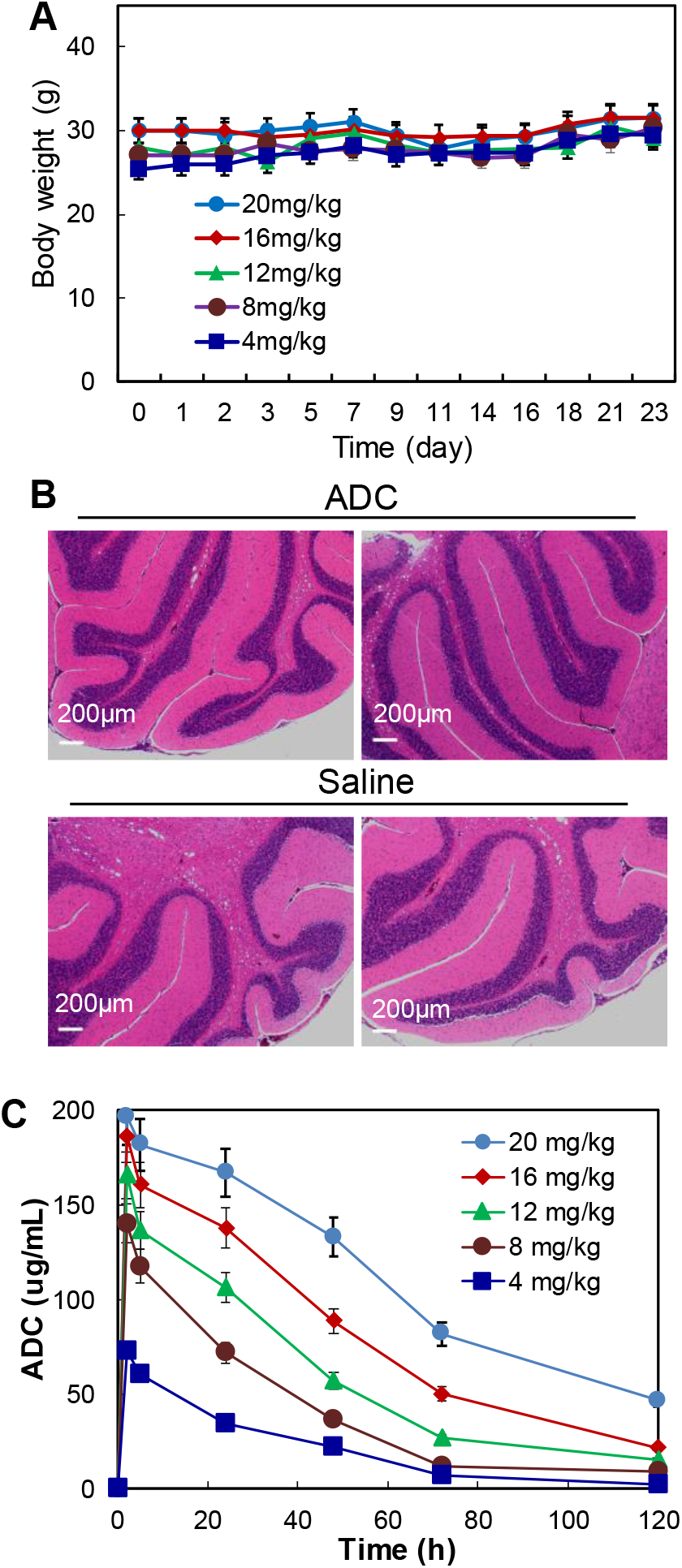
MTD and PK studies of ADC. **A,** MTD to test the effect of five ADC dosages including 4, 8, 12, 16 and 20 mg/kg-BW. **B,** H&E staining of brain tissues. Scale bar equals to 200 μm. **C,** PK to evaluate the stability and kinetics parameters of ADC (data represent mean ± SEM).

### Pharmacokinetics (PK)

The PK study was done by intravenously injecting ADC into s.c. NET xenografted mice at five different concentrations: 4, 8, 12, 16, and 20 mg/kg BW. Plasma samples were collected (10-50 μL each) from tail at time points of 0, 2, 8, and 16 hr, and 1, 2, 3, 5, and 7 days post-ADC injection and then titrated using an ELISA assay (Fig. 6C). The PK modeling demonstrates the recommended dose (D) = 3.78-14.30 mg/kg BW and recommended dosing interval (τ) = 4.40-9.10 days. Therefore, we selected a treatment dose of 8 mg/kg BW with administration interval of 4-5 days for the remaining *in vivo* anti-cancer study.

### *In vivo* anti-cancer efficacy

The mice bearing BON-Luc xenografts were treated in a dosing interval of 4.5 days with either anti-SSTR2 ADC (8 mg/kg), anti-SSTR2 mAb (8 mg/kg, control), or saline (vehicle, control) in three groups (*n* = 6). Fig. 7A showed that tumor growth was significantly inhibited with a tumor size reduction of 62-67% in ADC treatment group as compared with the control groups. The tumor fluorescence flux measured with IVIS showed 71-73% of growth reduction in treatment group compared to the control groups (Fig. 7B). The wet weight of the harvested tumors also confirmed the significant inhibition of tumor growth (Figs. 7C-D). There was no obvious difference among the three groups in overall body weight change (Fig. 7E). Western blotting analysis showed that SSTR2 expression was present in NET tumors during treatment (Fig. 7F). The surface staining of SSTR2 in tumors from ADC treatment group appeared to be lower than the control group (Fig. 7G), likely due to the NET cell death caused by ADC which was confirmed through H&E staining (Fig. 7H). This *in vivo* anti-cancer efficacy study demonstrates that the anti-SSTR2 mAb provides good drug delivery and the antibody-drug conjugate can effectively and safely inhibit NET growth.

**Figure 7.**
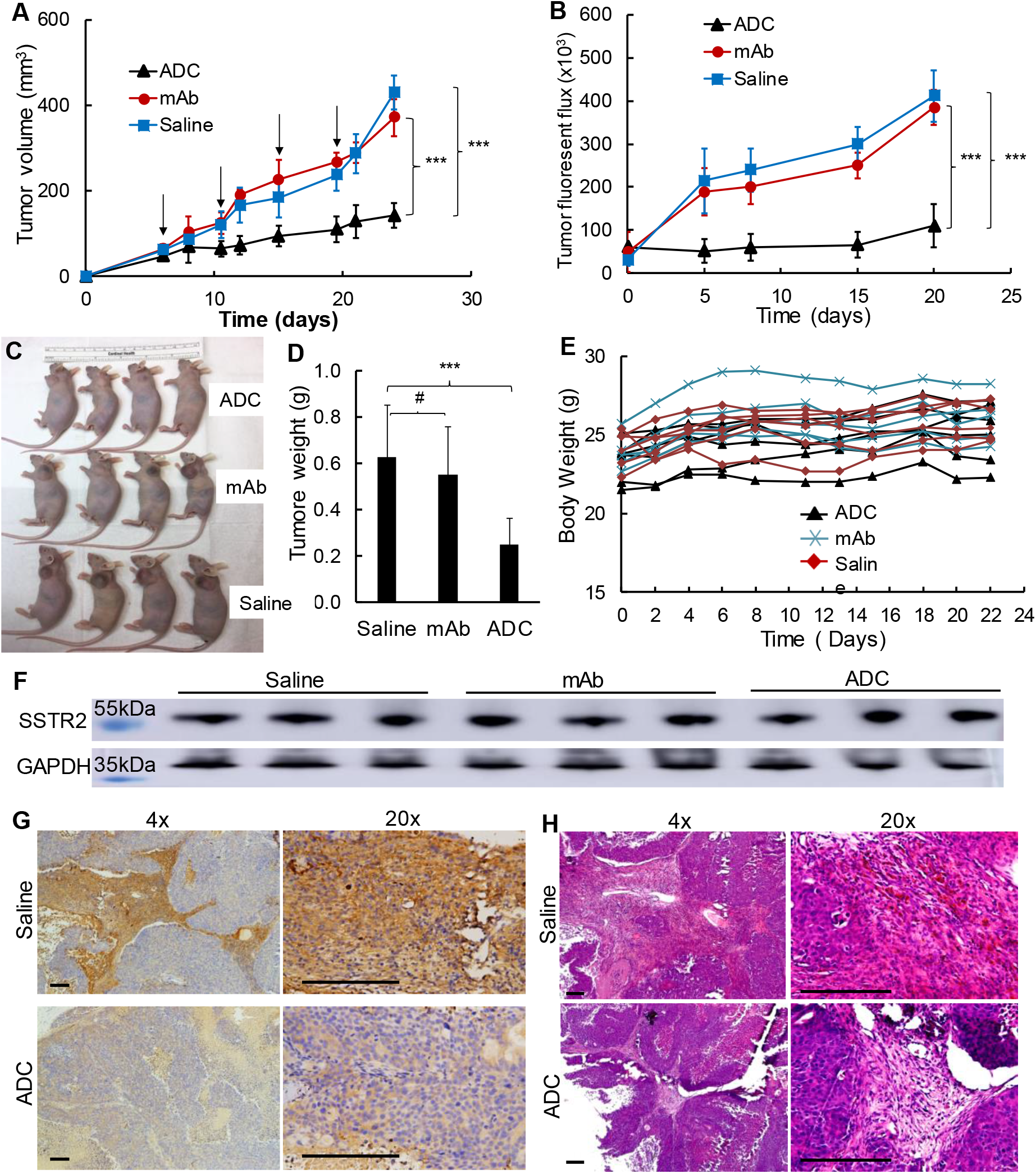
Anti-cancer efficacy study of ADC in NET (BON-Luc) xenografted mouse model. **A,** Tumor volume changes after BON-Luc cells injection and treatment (data represent mean ± SEM, *n* = 6). Tumor was measured with calipers, and calculated as ellipsoid. Black arrow indicating ADC (8 mg/kg BW) treatment date. **B,** Tumor fluorescence flux measurement with IVIS image system (data represent mean ± SEM, *n* = 6). **C,** Tumor bearing mice harvested. **D,** West weight of the tumors excised from harvested mice. **E,** Body weight of the mice during treatment. 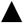: treatment group injected with ADC, 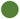: control group injected with mAb, and 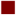: control group injected with saline. **F,** Western blotting of tumors from represented mice (*n* = 3). **G,** Anti-SSTR2 IHC staining of the saline and ADC treated tumors. **H,** H&E staining of saline or ADC treated tumor. Scale bar equals to 50 μm. *** *p* ≤ 0.001.

## Discussion

To develop effective and safe targeted cancer therapies, a unique biomarker that specifically defines the cancer cells from the non-cancerous cells must be identified and thoroughly characterized. The Human Atlas Project reported high mRNA expression of SSTR2 in several normal human tissues (such as brain), but the surface protein expression in these tissues (and other normal tissues) is low or undetectable. Although several studies reported SSTR2 protein expression in central nervous system, gastrointestinal tract, and pancreas,(44) the surface expression of SSTR2 in NET tissues is >20-fold higher than that in normal tissues. Considering ADC is a dose-dependent targeted therapy, the drastically high expression in NETs assures the safety to target SSTR2 and deliver therapeutic drugs. Moreover, our study and other studies(22–24) demonstrate that more than 70% of NET patients abundantly express SSTR2. All the results collected from patient tumor tissues, normal organs, and cell lines demonstrate that SSTR2 is an ideal receptor for targeted cancer therapy.

Similar to the finding in this study, literature shows that not all patients with NETs overexpress SSTR2.(45, 46) For example, it has been reported that only 45-66% of pulmonary NET patients and 80-95% gastroenteropancreatic NET patients overexpress SSTR2.(45) To benefit the SSTR2 negative patients, we have performed a comparative membrane proteomics study and found that the carcinoembryonic antigen-related cell adhesion molecule 1 (CEACAM1) has high expression in pancreatic NET cells (BON-1 and QGP-1) but not in non-NET cancerous pancreatic adenocarcinoma cells (PANC-1 and MiAPaCa-1) and non-cancerous fibroblast cell (WI-38). Literature has also reported high CEACAM1 expression in various cancers, including medullary thyroid cancer which represents a type of NET.(47, 48) Although further evaluation is needed, we can potentially use CEACAM1 as an alternative receptor of SSTR2 for some NET patients with minimal SSTR2 density.

In this study, we developed, characterized and confirmed a novel monoclonal antibody to target the identified SSTR2 receptor for NET therapy. Differently from the commercial mAb that was developed using the whole SSTR2 membrane protein as an immunogen, the anti-SSTR2 mAb was created using two extracellular domains of SSTR2 as immunogens in hybridoma technology. Therefore, our mAb showed a higher and more specific surface binding to NET cells than the commercial mAb. Furthermore, the developed new mAb can target both human and mouse SSTR2, so the exclusive accumulation of anti-SSTR2 mAb in human NET xenograft in the s.c. mouse model indicated the specific targeting in patients. Importantly, the maximum tolerated dose study did not detect any body weight or behavior changes of the mice treated with dose of up to 20 mg ADC/kg BW. The H&E staining on murine brain tissue which has the highest mRNA expression of SSTR2 did not show any evidence of cellular damage or morphology change. Altogether, the developed anti-SSTR2 mAb can specifically target the SSTR2-overexpressing NET cell lines, patient-derived tissues and xenografts. Therefore, it is evident that the new mAb has the great potential to specifically deliver highly potent small molecules to NET.

For the first time, we developed a SSTR2-targeted therapy for NET treatment in the form of monoclonal antibody-drug conjugate. Our novel ADC demonstrates higher therapeutic values than other therapies. For instance, the tumor growth in s.c. xenograft mice is significantly reduced upon treatment with anti-SSTR2 ADC, which is more effective than the chemotherapies (e.g., Everolimus, Sunitinib, Octreotide, and Lanreotide) under investigation and radionuclide therapy (i.e. Lutathera) approved by FDA. Moreover, ADC has multiple advantages compared to other therapies, including the enhanced cellular uptake via strong surface binding, high cytotoxicity of the delivered small molecule payload, and minimal side effects.

In addition to the NET-specific targeting of anti-SSTR2 mAb and the high potency of conjugated drug, we also found that the mAb could downregulate the cell proliferation associated PI3K/AKT signaling, downregulate the oncogene cyclin D1, and upregulate the cell cycle associated p21. These findings indicated SSTR2-targeting ADC could serve as a novel multi-purpose biologic with clinical potentials to directly cause cell death by releasing a cytotoxic payload and inhibite tumor cell growth via the SSTR2-mediated modulation of signaling cascades. Other studies have also reported multiple mechanisms that could drive anti-tumor effects mediated by SSTR2, such as apoptosis, regulation of cyclin-dependent kinase inhibitors, and inhibition of proliferation signaling.(49, 50) It is necessary to further investigate the synergy of anti-SSTR2 mAb and ADC for NET treatment *in vivo* using a sporadic MTC mouse model or humanized mouse model in future.

In conclusion, our anti-SSTR2 ADC has higher therapeutic values than traditional chemotherapy, radiotherapy, and surgery to treat NE cancers due to its capability or potential to: 1) target and treat the metastatic nodules; 2) reduce undesirable side effects; and 3) effectively reduce NE cancer growth. Similar to other FDA approved receptors, SSTR2 is not an absolute NET-specific receptor, so it is imperative to further evaluate the potential side effects. The combination of the facts that SSTR2 expression in NETs is greater than normal tissues, SSTR2 has little or undetectable surface expression in normal organs, and ADC is a dose-dependent treatment strategy could minimize the possible off-target side effects. Integrated with other therapies, the targeted therapy developed in this study has great potential to improve the quality of life and survival rate of patients with NE cancers.

## Data Availability Statement

The data that support the findings of this study are available from the corresponding author upon reasonable request.

